# Decarboxylation of indoleacetic acid illuminates its evolution into a plant hormone

**DOI:** 10.64898/2026.09.07.749952

**Authors:** Roman Skokan, Vojtěch Schmidt, Petre I. Dobrev, Vojtěch Knirsch, Karel Müller, Barbora Svobodová, Jozef Lacek, Petr Maršík, Stanislav Vosolsobě, Yoan Coudert, Jan Petrášek

## Abstract

The evolution of indoleacetic acid (IAA) into the plant hormone auxin remains unclear. Here, we compared IAA response and metabolism in land plants and their closest living relatives, the streptophyte algae, using chemical treatments, phytohormone profiling, and targeted and untargeted metabolomics. Streptophyte algae showed neither plant-like auxin responses nor canonical IAA metabolism. Here, we demonstrate IAA side-chain decarboxylation as a novel pathway of cellular IAA metabolism, operating in streptophyte algae and land plants alike. Decarboxylation also underpinned IAA’s innate photolability and yielded cytotoxic intermediates, which also accumulated endogenously under simulated sunlight. Together, these findings establish IAA decarboxylation as a major, conserved metabolic pathway, and suggest that the metabolic constraints arising from IAA’s instability and indirect cytotoxicity contributed to its adoption as a plant hormone.

## Main text

Indole-3-acetic acid (IAA) is a widespread biological molecule, known as auxin for its morphogenic influence in land plants (Schmidt et al., 2024; Vanneste et al., 2025). The canonical molecular mechanisms of auxin signaling, biosynthesis and transport, first described in angiosperms, are conserved across land plants (Carrillo-Carrasco et al., 2023). Their closest living relatives, the streptophyte green algae, lack genomic signatures of plant-like auxin biosynthesis and signaling, but show independent functional and molecular evidence of cellular IAA transport (Boot et al., 2012; Skokan et al., 2019; Bowman et al., 2021). Physiological response to applied auxins has been individually tested in multiple species of streptophyte algae, but the results lack common consensus (Wood & Berliner, 1979; Dibb-Fuller & Morris, 1992; Klämbt et al., 1992; Jin et al., 2008; Kwiatkowska et al., 2014; Ohtaka et al., 2017; Kuhn et al., 2024; Zeng et al., 2024; Carrillo-Carrasco et al., 2025; Kurtović et al., 2025). In this study, we aim to elucidate why IAA had been adopted as a prominent morphogen during plant terrestrialization. We do this by searching for remnant links in IAA homeostasis and physiological influence between land plants and streptophyte algae. With insights gained from auxin treatments and IAA metabolism, we offer our hypothesis on what had made IAA a suitable candidate to evolve into a signaling molecule.

### Auxin treatments

Land plants show growth responses to applied auxin at nanomolar doses, and severe phenotypes manifest at units of micromol. We performed an analogous analysis in four selected strains of streptophyte algae spanning three separate lineages: *Klebsormidium nitens* (Klebsormidiophyceae), *Coleochaete scutata* (Coleochaetophyceae), *Spirogyra pratensis* and *Closterium peracerosum-strigosum-littorale* complex (both Zygnematophyceae), further referred to by generic names (**Fig. 1A**). These algae were exposed to a range of concentrations (0.01 to 100 µM, in 10^1^ increments) of IAA, 1-naphthaleneacetic acid (1-NAA) and 2-NAA. These represent a native plant auxin, a synthetic auxin and a close structural analog without auxin activity, respectively (Dahlke et al., 2009; Paponov et al., 2019). Culture growth was observed and quantified 7-10 days post inoculation (**Fig. 1B,C; Table S1**). 100 µM IAA proved to be a generally detrimental dose, ranging from mild culture growth inhibition *(Klebsormidium)* to lethality (Zygnematophyceae). 10 µM IAA provoked a comparatively milder reaction which differed by species. Any lower dose had no significant effect on culture growth. Importantly, there was a general overlap in the influence of IAA, 1-NAA and 2-NAA, except in *Coleochaete* (**Fig. 1C**). Our results stand in contrast to an earlier study in the desmid *Micrasterias*, where ca. 2 µM IAA promoted culture growth (Wood & Berliner, 1979). However, we concur with more recent research on *Klebsormidium* and *Penium* regarding the high effective doses of applied IAA and the overlapping influence of similar chemical compounds (Ohtaka et al., 2017; Carrillo-Carrasco et al., 2025). In contrast to land plants, streptophyte algae showed no dose-dependent, biphasic modulation of growth; rather, elevated concentrations of IAA and structurally related compounds were toxic. These results indicate that IAA lacks auxin-like signaling function in streptophyte algae, in line with the established evolution of auxin signaling (Hernandez-Garcia & Weijers, 2026).

**Figure 1.**
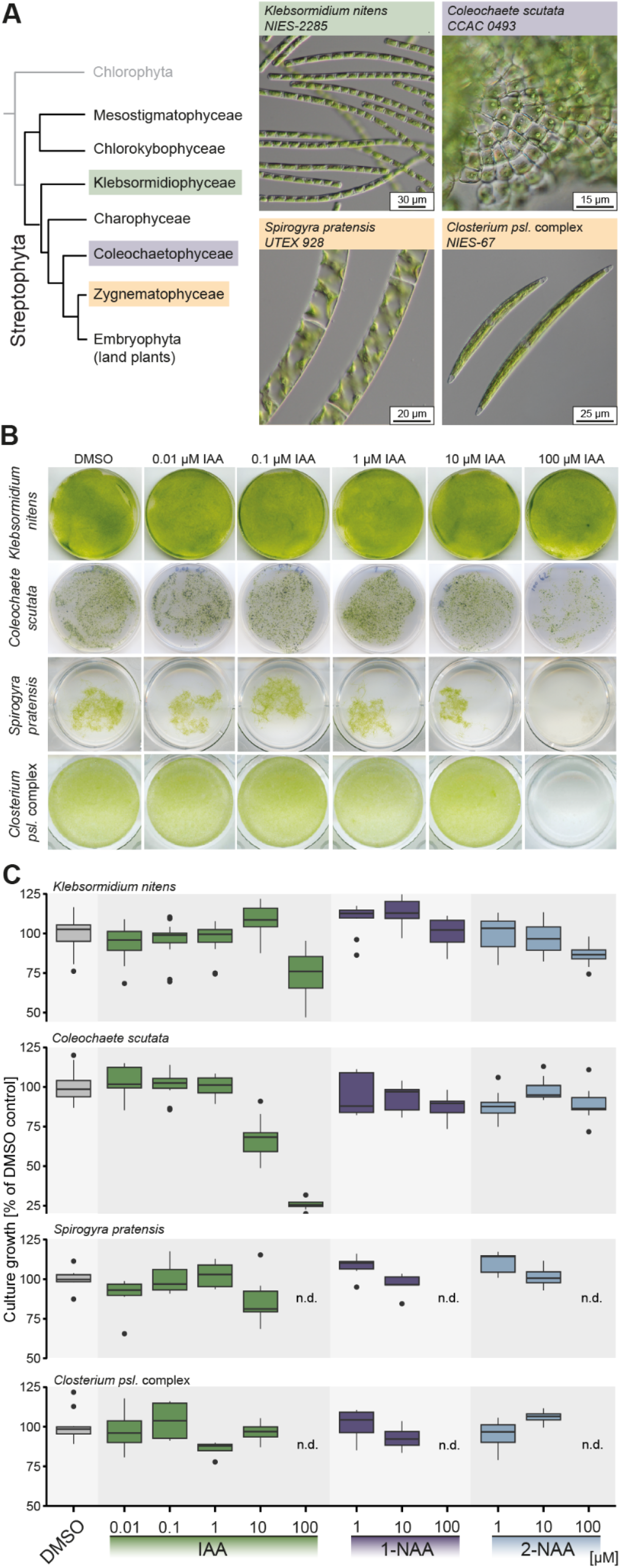
**(A)** Relationship of selected charophyte representatives used in this study within Streptophyta with DIC images; scale bar = 50 μm. (**B)** Culture growth response of charophytes to applied auxins; the effect of IAA. 14 days cultivation. (**C)** Culture growth response of charophytes to applied auxins. 14 days of cultivation. Indole-3-acetic acid, IAA; 1-naphthaleneacetic acid, 1-NAA; 2-naphthaleneacetic acid, 2-NAA. Culture growth as fold change vs. moc (DMSO) treatment, fresh weight.

### Metabolism of applied IAA in algae and plant cells: Unique patterns and a novel pathway

In angiosperms, the auxin action of IAA is coupled with its meticulous intracellular homeostasis (Casanova-Sáez et al., 2021; Hayashi et al., 2021). To gain comparison in streptophyte algae, we supplemented the culture media of four algal species with [^13^C_6_]IAA and traced its conversion into downstream compounds over 24 hours. [^13^C_6_]IAA was taken up and overall gradually depleted in the algal biomass (**Fig. 2A)**, with a corresponding drop in the culture medium, but not in a control medium without biomass (**Fig. S1**). This attests to an active uptake and turnover of [^13^C_6_]IAA by streptophyte algae. However, [^13^C_6_]IAA metabolism in streptophyte algae differed from the pattern known from land plants. While seed plants have been known to process applied IAA mainly through conjugation to amino acids and glucose, very little such metabolization of [^13^C_6_]IAA was detected in streptophyte algae (**Fig. 2B**). Like land plants, streptophyte algae oxidized [^13^C_6_]IAA at the indole core (**Fig. 2C**). However, [^13^C_6_]-2-oxo-IAA was not accumulated over time, as previously reported in the moss *Physcomitrium patens* (Abitbol-Spangaro et al., 2025). Moreover, angiosperm-like oxidation of conjugated IAA was not detected in streptophyte algae (**Fig. 2B**), in agreement with the lack of non-oxidized conjugates. Although our study concurs with earlier reports that streptophyte algae can produce marginal amounts of angiosperm-like IAA conjugates, it is clear that the metabolisation of supplemented [^13^C_6_]IAA proceeds mainly through other means in the algae.

**Figure 2.**
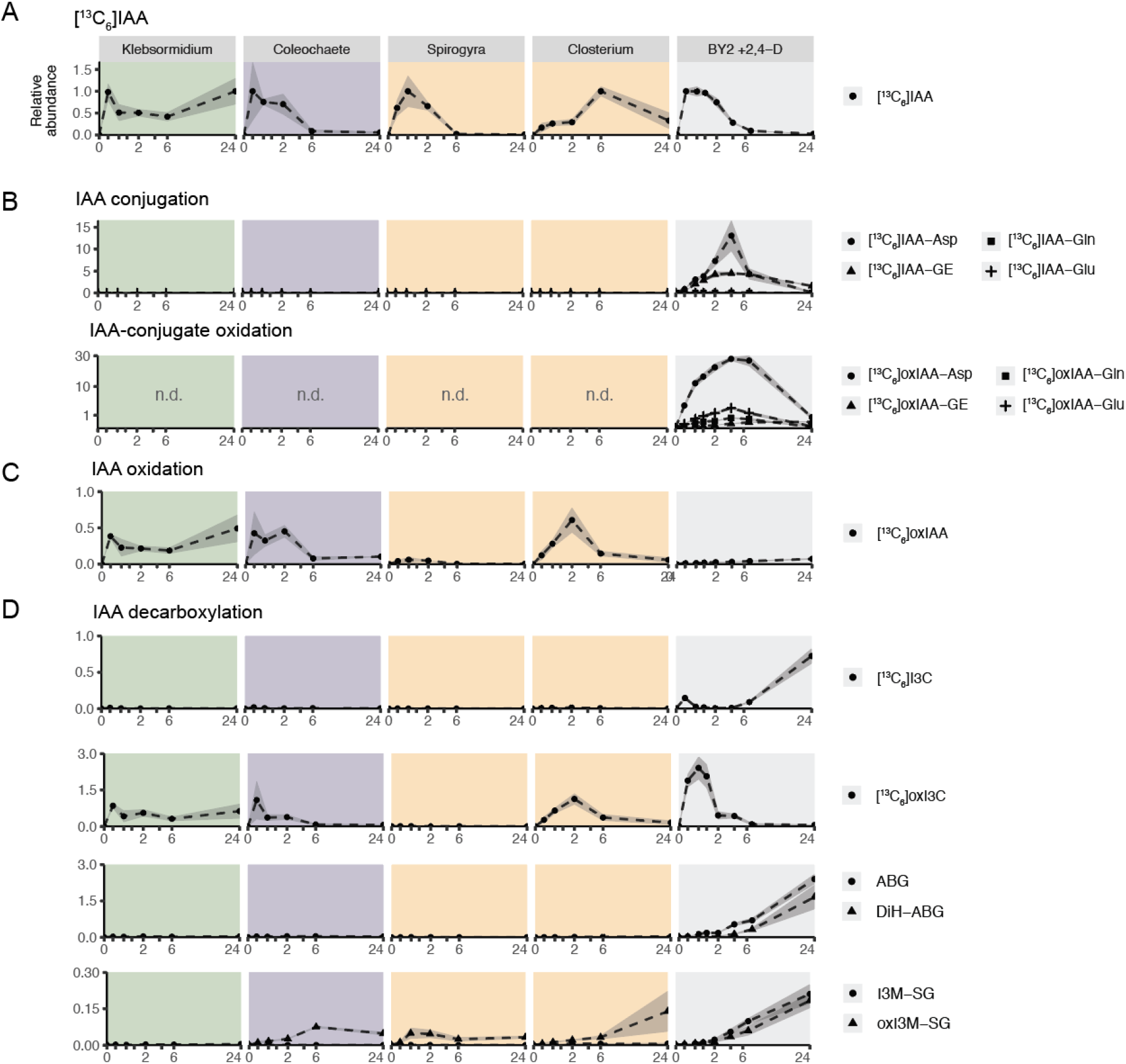
[^13^C_6_]IAA metabolism over 24 hours in four streptophyte algae and BY-2. **(A)** [^13^C_6_]IAA. **(B)** [^13^C_6_]IAA conjugation to glucose and amino acids, and the oxidation of these compounds. **(C)** [^13^C_6_]IAA oxidation. **(D)** [^13^C_6_]IAA decarboxylation. Abbreviations: IAA, indole-3-acetic acid; oxIAA, 2-oxo-indole-3-acetic acid; IAA-Asp, IAA-aspartate; IAA-Glu, IAA-glutamate; IAA-GE, IAA-glucosyl ester; oxIAA-GE, oxIAA-glucosyl ester; oxIAA-Glu, oxIAA-glutamata; oxIAA-Gln, oxIAA-glutamine; I3C, indole-3-carbinol; I3C-GSH, I3C-glutathione; ABG, ascorbigen; DiHABG, dihydroascorbigen, oxI3C, 2-oxo-indole-3-carbinol; Mene-oxI, 3-methylene-2-oxindole; oxI3C-GSH, oxI3C-glutathione; n.d., not detected.

Seeking to explain [^13^C_6_]IAA metabolization in green algae, we re-assessed the known spectrum of IAA metabolism in the green lineage. In fact, historical research indicates that even angiosperms may possess a richer IAA metabolic spectrum than is currently accepted, indicating the possible existence of unknown pathways of IAA metabolism. Recently, IAA decarboxylation had been described as an endogenous IAA catabolic pathway in the Bright Yellow-2 (BY-2) cell culture of *Nicotiana tabacum* (Dobrev et al., 2023), an established model to study cellular auxin homeostasis (Müller et al., 2021), and performed an untargeted analysis of IAA metabolism. The established pathways of IAA conjugation and oxidation were confirmed in BY-2, but additional peaks corresponding to unknown IAA metabolites were also detected (**Fig. S2**) and subsequently identified as the products of IAA decarboxylation at the side chain (**Fig. S3**). To determine the evolutionary extent of IAA decarboxylation, we updated our detection method and re-examined the previous [^13^C_6_]IAA metabolic assays in other organisms. We found that all four streptophyte algae processed [^13^C_6_]IAA by decarboxylation, with the same molecular products as in BY-2 (**Fig.2D**). Moreover, [^13^C_6_]IAA decarboxylation was also detected in the model bryophyte *Physcomitrium patens* (dataset produced for Abitbol-Spangaro et al., 2025) (**Fig. S4**). Additional experiments in BY-2 have established individual steps within the IAA decarboxylation pathway (**Fig. S5**) and confirmed its enzymatic nature (**Fig. S6**). Insofar, IAA decarboxylation has been known mainly as an *in vitro* phenomenon, with indirect indications in living plant material through feeding of labelled IAA and subsequent detection of released CO_2_. Here, we firmly established the existence of this pathway in living cells of land plants and streptophyte algae.

### IAA decarboxylation: A major endogenous pathway

So far, our experiments have dealt with IAA applied externally and in overabundance, compared to naturally occurring levels in algal and plant tissues (Schmidt et al., 2024). To assess the relevance of IAA decarboxylation in a native context, we performed phytohormone profiling in two species of angiosperms (*Arabidopsis thaliana, Nicotiana benthamiana*), two bryophytes (*Marchantia polymorpha, P. patens*) and the above-listed four streptophyte algae. In agreement with [^13^C_6_]IAA feeding assays, conjugated IAA and oxIAA were detected in the two angiosperms, whereas little to none were found in bryophytes and streptophyte algae (**Fig. 3**). Oxidized IAA was present in all tested organisms (**Fig. 3**). Finally, IAA decarboxylation was positively identified in all land plants and streptophyte algae (**Fig. 3**). In fact, the products of IAA decarboxylation dominated the total pool of endogenous IAA metabolites screened in this study, across all tested species (**Fig. 3**). Therefore, the insofar neglected IAA decarboxylation represents a major route of endogenous IAA metabolism in both land plants and streptophyte green algae. Notably, *A. thaliana* does not represent a suitable model organism to study IAA decarboxylation due to the interference of the *Brassicaceae*-specific glucosinolate metabolism, which likewise produces indole-3-carbinole (Katz et al., 2015).

**Figure 3.**
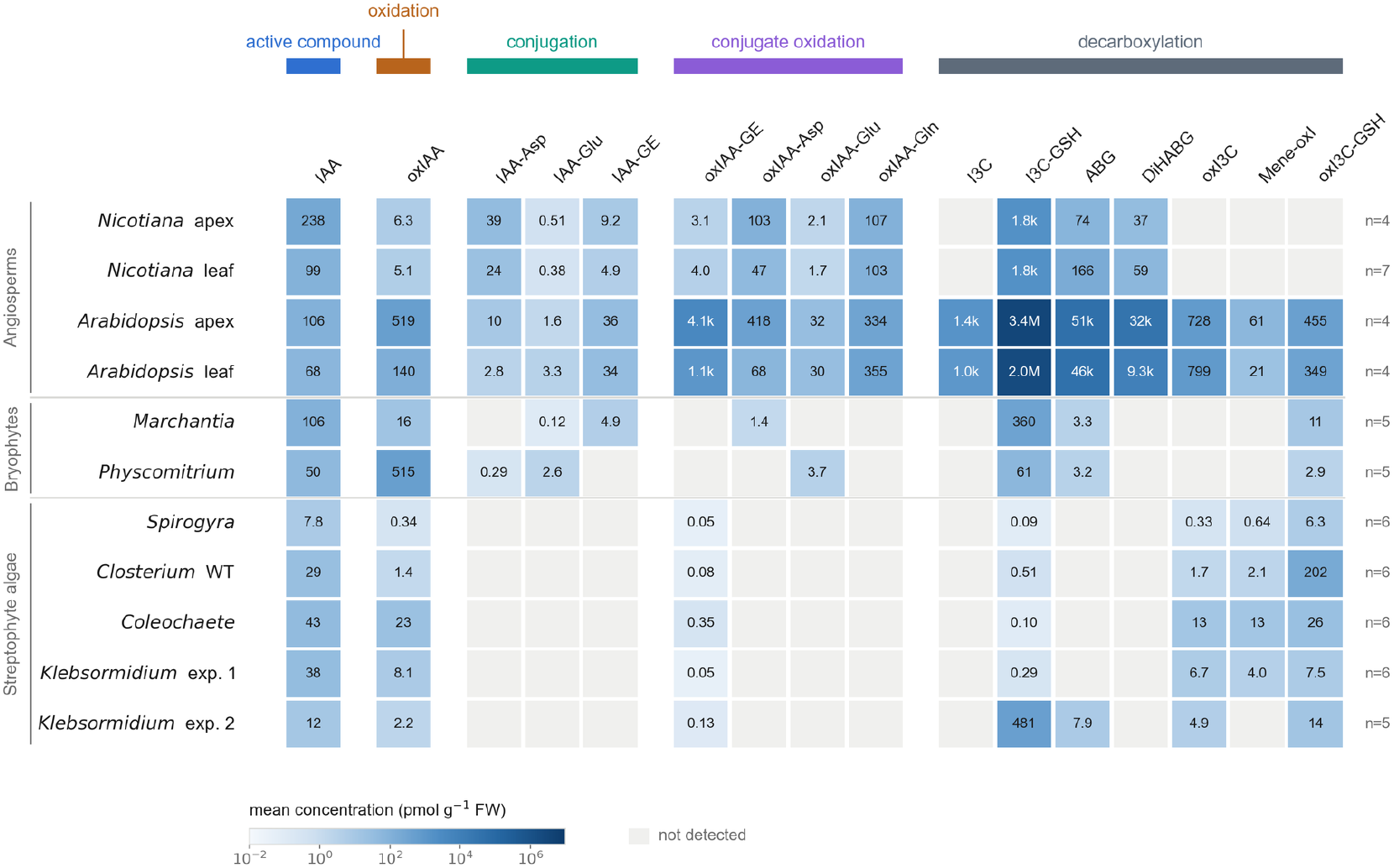
Endogenous content of IAA and its metabolites in land plants and streptophyte algae under standard culture conditions.

### Enzymatic vs. non-enzymatic modes of IAA decarboxylation

In the [^13^C_6_]IAA feeding experiment of streptophyte algae, no metabolic changes were detected in the pure medium control (**Fig. S1**). In BY-2, the culture filtrate did induce IAA decarboxylation, but not after boiling (**Fig. S6**). These observations indicate that IAA decarboxylation proceeded enzymatically and, to an extent, in the apoplast. However, IAA decarboxylation, coupled with simultaneous side chain oxidation, has been known to also proceed non-enzymatically, mainly due to the influence of certain light wavelengths and chemical components of the culture media (Stasinopoulos & Hangarter, 1990; Brennan, 1996). Therefore, we tested the stability of IAA supplemented into the culture medium under our culture and experimental conditions, in a cell-free setup. We found IAA to be stable over two weeks under the conditions corresponding to algal culture, auxin treatments and metabolic assays (**Fig. 4A, Fig. S1**). However, a slight alteration in the culture setup, specifically the inclusion of a marginal amount of UV light (**Fig. S7**), resulted in considerable IAA degradation over the same period (**Fig. 4A**). Remarkably, this IAA breakdown included indole ring oxidation and decarboxylation (**Fig. 4A**), i.e. largely overlapped with active [^13^C_6_]IAA metabolism in streptophyte algae. We further inquired into the molecular stability of IAA, using 4 hour incubation under stronger illumination mimicking natural sunlight (**Fig. S7**). Thus, we revealed that IAA shares its photolability with similar compounds based on the indole core, namely indole-3-carboxylic acid and indole-3-propionic acid (**Fig. S7**). Of note, the latter two compounds are not natively produced in land plants or streptophyte algae. However, the native but non-indolic compound phenylacetic acid (PAA), which is known as a weak auxin (Cook, 2019), was stable under the same conditions (**Fig. S7**). Therefore, the innate instability of IAA is derived from its indole core, as indicated by earlier research using other indolic compounds. Strikingly, the abiotic photooxidation of IAA produces certain identical compounds as its enzymatic metabolism, as measured in streptophyte algae and land plants.

**Figure 4.**
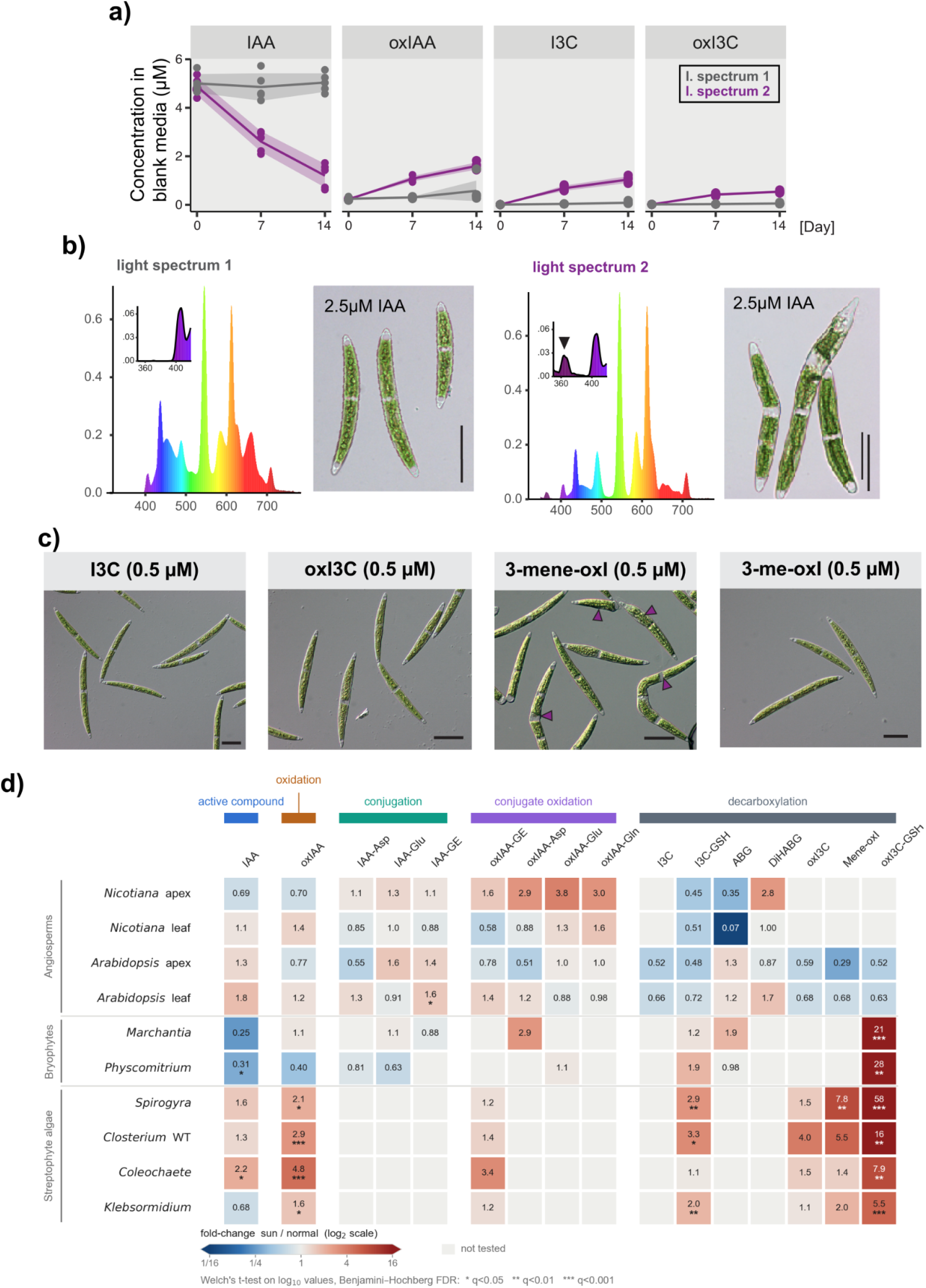
**(a)** IAA decay in C medium within glass Erlenmeyer flasks under different illumination. Light spectra correspond to (b). **(b)** Phenotype of IAA-treated *Closterium* cells (14 days) cultured under different illumination conditions. **(c)** Individual treatments of *Closterium* cells (3 days) with products of IAA decarboxylation.I3C, indole-3-carbinol; oxI3C, oxindole-3-carbinol; 3-mene-oxI, 3-methylene-2-oxoindole; 3-me-oxI, 3-methyl-2-oxoindole. **(d)** Endogenous content of IAA and its metabolites in land plants and streptophyte algae, 4-hour incubation under simulated sunlight vs. standard culture conditions (fold change).

### Biological impacts of IAA decarboxylation

In both land plants and streptophyte algae, the products of IAA decarboxylation were ultimately conjugated to nucleophilic scavengers such as glutathione and ascorbate, indicating their reactive nature (**Fig. 2D**). Indeed, cytotoxic compounds have been known to result from acellular oxidative decarboxylation of supplemented IAA. In contrast to a relatively mild growth reaction of streptophyte algae to IAA treatment under confirmed, IAA-stable conditions (**Fig. 1C, Fig. S7**), the inclusion of a marginal amount of UV light (**Fig. 4A**) brought about dramatic culture growth inhibition coupled with cell malformations in *Closterium* at 2.5 µM of added IAA (**Fig. 4B**). Other tested compounds including 1-NAA or PAA did not induce such effects (**Fig. S8**). Expecting that a reactive product of UV-induced IAA breakdown might be responsible, we purified stocks of compounds resulting from IAA decarboxylation and performed individual, 3-day treatments of *Closterium* under IAA-stable conditions. The early products of IAA decarboxylation, namely indole-3-carbinol (I3C) and 2-oxo-indole-3-carbinol (oxI3C), did not alter *Closterium* cell morphology (**Fig. 4C**). By contrast, 3-methylene-2-oxo-indole (3-mene-oxI), a downstream product of oxI3C, replicated the previously observed *Closterium* cell malformations, specifically due to its reactive methylene group (**Fig. 4C**). These observations concur with previous reports of the physiological effects of IAA decarboxylation products, mainly 3-mene-OxI, in bacteria, yeast and animal cells (Fukuyama & Moyed, 1964; Folkes & Wardman, 2003; Papagiannakis et al., 2017). Therefore, we confirmed the cytotoxicity of at least one product of IAA decarboxylation, capable of adversely affecting algal growth and cell morphology when the capacity for its quenching is overwhelmed.

Finally, we speculated whether the susceptibility of IAA to photooxidative breakdown could be physiologically relevant under natural conditions such as midday sun, where land plants and streptophyte algae face high illumination and oxidative stress. Therefore, we exposed two angiosperms, two bryophytes and four streptophyte algae to four hours of simulated sunlight (**Fig. S7**) and assessed their endogenous IAA profiles. Compared to standard culture conditions, the young leaves of ‘sunlit’ angiosperms did not show significant changes in IAA metabolism (**Fig. 4D**). By contrast, the artificial sunlight caused a considerable upsurge of IAA decarboxylation in bryophytes and especially streptophyte algae. In particular, we detected a considerable accumulation of the glutathione-bound version of 3-mene-oxI (oxI3M-SG) (**Fig. 4D**); the same compound implicated in poisoning *Closterium* cells. In total, while endogenous IAA was protected from photooxidative breakdown in young angiosperm leaves under simulated sunlight, bryophytes and streptophyte algae suffered a surge in IAA decarboxylation, incurring the metabolic cost of quenching its cytotoxic intermediates.

## Conclusion

The accepted model of IAA metabolism, derived mainly from angiosperms, has fallen short of providing a full picture. Here, we establish IAA decarboxylation as one of the major pathways of IAA metabolism, originating well before plant terrestrialization and the evolution of known auxin signaling (Hernandez-Garcia & Weijers, 2026). This cellular decarboxylation of IAA is analogous to its non-enzymatic, photooxidative breakdown, described in earlier *in vitro* research. Accordingly, IAA decarboxylation was promoted in living cells exposed to simulated sunlight. We propose the following scenario of IAA’s evolution into auxin in the streptophyte algal ancestor of land plants: IAA’s molecular instability, coupled with non-specific cellular enzymes, facilitated signal attenuation, while the reactivity of its decarboxylation products drove the evolution of dedicated IAA homeostasis. The ancestral physiological role would be exerted either by IAA itself or by its reactive catabolites, as has been proposed for certain growth processes even in angiosperms (Bhattacharyya et al., 1986; Tuli, 2000). Conceivably, a hypothetical ancestral function of IAA in morphologically simple organisms would exploit IAA’s instability in an extracellular environment: exported by the deeply conserved auxin transport protein families (Vosolsobě et al., 2020), IAA would be readily degraded by sunlight and cell wall peroxidases. IAA itself or its reactive catabolite(s) would act as an unstable signal, or even as a form of antimicrobial repellent. In the framework of this study, IAA’s ultimate evolution into auxin began with managing its own chemical liability.

## Materials & Methods

### Chemicals

All chemicals and reagents were obtained from Merck/Sigma Aldrich (Darmstadt, Germany), unless stated otherwise.

### Algal and plant strains and their cultivation

*Closterium peracerosum-strigosum-littorale* complex (NIES-67 and NIES-68) and *Klebsormidium nitens* (NIES-2285) were obtained from Microbial culture collection of National Institute for Environmental Studies (NIES; Tsukuba, Japan), *Spirogyra pratensis* (UTEX 928) was obtained from the Culture Collection of Algae, University of Texas (UTEX; Austin, USA), and *Coleochaete scutata* (CCAC 0493) was obtained from Central Collection of Algal Cultures, Duisburg University (CCAC; Duisburg, Germany). *Closterium psl*. complex was cultured in liquid C medium, while *Klebsormidium nitens* and *Coleochaete scutata* were cultured on solid C medium (1.5% agar; w/v). *Spirogyra pratensis* was grown in liquid Bold’s Basal Medium (BBM, Merck B5285). All media were supplemented with the following vitamins: B_1_, B_12_ (10 mg/l each), B_7_ (2 mg/l), which were added from individual 1000x concentrated stocks (prepared in dH_2_O, sterilized via 0.22µm filtration; Millex SLGS033SS) into cooled agar media before solidification, or directly into liquid media prior to inoculation. For regular propagation, liquid cultures were supplemented with soil extract. Soil extract was prepared by mixing soil with dH_2_O in a 1:3 ratio (v/v), followed by sterilization via autoclaving. The soil, free of noticeable leaf litter, was collected in a submontane forest of European beech (*Fagus sylvatica; 50*.*82722N, 14*.*46689E*). The suspension was then filtered through filter paper overnight and autoclaved twice more. Finally, soil extract was passed through a 0.22µm filter (Millex SLGS033SS), stored at −20°C, and added into media prior to inoculation. Algal strains were subcultured every 6-8 weeks and cultivated at 23°C, 16:8 hours light:dark regime, illuminated at 30 µE light intensity using the combination of two fluorescent light tubes: Osram Fluora TLD 36W (Osram Licht AG, Germany) and Philips Master TLD Super 36W (Koninklijke Philips N.V., The Netherlands).

### Chemical treatments

For treatments, charophyte strains were inoculated into or onto culture media (soil extract omitted), and pH adjusted to 5.5. IAA, 1-NAA, 2-NAA, I3C, and 3-me-oxI were dissolved in DMSO to prepare concentrated stocks. oxI3C and 3-mene-oxI were dissolved in 5% and 40% acetonitrile (ACN; v/v), respectively, reflecting the solvent conditions used for their HPLC-based purification (see below). The equivalent volumes of respective solvents were used as mock controls.

### Feeding of algae with [^13^C_6_]IAA stable isotope

Two-week-old cultures of *Coleochaete scutata, Spirogyra pratensis, Closterium psl*. complex, and 1-week-old culture of *Klebsormidium nitens* were harvested and incubated in 50 ml Falcon tubes (each strain in three biological replicates from independently grown cultures) containing 6.5 ml of fresh, liquid C medium. The medium was modified by omitting TRIS and vitamins, and adjusting the pH to 5.5. Medium was supplemented with 1µM [^13^C_6_]IAA stable isotope (Cambridge Isotope Laboratories, Tewksbury, MA, USA). Biomass was collected from each tube in two technical replicates at specific timepoints (10 min, and 0.5, 2, 6, 24 hours) by pipetting, placing on a 20 µm nylon net filter (Merck Millipore, NY2004700), washing by dH_2_O to remove any residual medium while draining using underpressure. Biomass was promptly re-weighed, flash frozen in liquid nitrogen, and stored at −80°C. Corresponding media samples were sampled at T0 and 24h per two replicates (100 µl each) by pipetting.

### LC-MS analysis

Biomass samples (10-50 mg, fresh weight) were homogenized with 1.5mm zirconium beads using a FastPrep-24 instrument (MP Biomedicals, CA, USA) with 100 µl of 1M HCOOH in water. 100 µl of 1M HCOOH was also added to liquid media samples (100 µl, not homogenized). After centrifugation at 30,000 x g for 20 minutes at 4°C and re-extraction, the combined supernatant was applied to an SPE Oasis HLB 10mg 96-well plate (Waters, Milford, MA, USA). The supernatant was pushed through the SPE plate using a Pressure+96 manifold (Biotage, Uppsala, Sweden). The 96-well SPE plate was washed three times with 100 µl of dH_2_O. Samples were eluted with 100 µl 50% acetonitrile/water (v/v). An aliquot of the SPE eluate was injected into the LC-MS system. Metabolites were separated on a Kinetex EVO C18 column (2.6 µm, 150 x 2.1 mm, Phenomenex, Torrance, CA, USA). The mobile phases consisted of A) 5 mM ammonium acetate and 2 µM Medronic acid in water and B) 95/5 acetonitrile/water (v/v). The following gradient program was used: 5% B at 0 min, 7% B at 0.1 to 5 min, 10 to 35% at 5.1 to 12 min, 100% B at 13 to 14 min, and 5% B at 14.1 min. Analysis was performed on an LC-MS system consisting of a UHPLC 1290 Infinity II (Agilent, Santa Clara, CA, USA) coupled to a 6495 Triple Quadrupole mass spectrometer (Agilent). MS analysis was performed in MRM mode. The following MRM transitions were used: [^13^C_6_]IAA (-) 180>136; [^13^C_6_]oxIAA (+) 198>152. Data acquisition and processing were performed using Mass Hunter software B.08 (Agilent).

### Preparation of IAA decarboxylation metabolites

I3C and 3-me-oxI were purchased from Merck. oxI3C was synthesized by photooxidation of IAA in the presence of riboflavin according to (Hope and Ordin 1971), followed by HPLC purification avoiding sample preconcentration. Purification was done using HPLC pump, diode array detector (Ultimate 3000, Thermo-Fisher Scientific, Waltham, MA, USA) and fraction collector FC 203B (Gilson, Middleton, WI, USA). Separation was performed on a Kinetex C18 column (5 µm, 150 x 4.6 mm, Phenomenex) at a flow rate of 0.6ml/min. The mobile phases consisted of A) water, and B) 95/5 acetonitrile/water (v/v). The linear gradient program was: 5-45% B for 20 min, 45-95% B for 1 min, 95 % B for 1 min, 95-5% B for 1 min, and 5% B for 5 min. 3-methylene-2-oxindole was produced from oxI3C by incubating it in an acidified buffer solution (100 mM ammonium acetate in water, pH 4) at room temperature for one day, followed by HPLC purification as described above.

## Notes

### Competing Interest Statement

The authors have declared no competing interest.

### Funding

- European Regional Development Fund programme Johannes Amos Comenius: CZ.02.01.01/00/22_008/0004581

## References

1. Abitbol-Spangaro, J. et al. Robust branch patterning in moss shoots via symplasmic auxin diffusion. Current Biology 35, 5238–5251.e10 (2025).

2. Bhattacharyya, R. N., Chattopadhyay, K. K. & Basu, P. S. Auxin activity of 3-hydroxymethyl oxindole and 3-methylene oxindole in oat. Biol Plant 28, 219–226 (1986).

3. Boot, K. J. M., Libbenga, K. R., Hille, S. C., Offringa, R. & Van Duijn, B. Polar auxin transport: an early invention. Journal of Experimental Botany 63, 4213–4218 (2012).

4. Bowman, J. L., Flores Sandoval, E. & Kato, H. On the Evolutionary Origins of Land Plant Auxin Biology. Cold Spring Harb Perspect Biol 13, a040048 (2021).

5. Brennan, T. M. Decarboxylation of lndole-3-Acetic Acid and Inhibition of Growth in Avena sativa Seedlings by Plant-Derived Photosensitizers*. Photochem & Photobiology 64, 1001–1006 (1996).

6. Carrillo-Carrasco, V. P. et al. Auxin and tryptophan trigger common responses in the streptophyte alga Penium margaritaceum. Current Biology 35, 2078–2087.e4 (2025).

7. Carrillo-Carrasco, V. P., Hernandez-Garcia, J., Mutte, S. K. & Weijers, D. The birth of a giant: evolutionary insights into the origin of auxin responses in plants. EMBO J 42, EMBJ2022113018 (2023).

8. Casanova-Sáez, R., Mateo-Bonmatí, E. & Ljung, K. Auxin Metabolism in Plants. Cold Spring Harb Perspect Biol 13, a039867 (2021).

9. Cook, S. D. An Historical Review of Phenylacetic Acid. Plant and Cell Physiology 60, 243–254 (2019).

10. Dahlke, R. I., Lüthen, H. & Steffens, B. The auxin-binding pocket of auxin-binding protein 1 comprises the highly conserved boxes a and c. Planta 230, 917–924 (2009).

11. Dibb-Fuller, JenniferE. & Morris, DavidA. Studies on the evolution of auxin carriers and phytotropin receptors: Transmembrane auxin transport in unicellular and multicellular Chlorophyta. Planta 186, (1992).

12. Dobrev, P. I. et al. Study of auxin metabolism using stable isotope labeling and LCMS; evidence for in planta auxin decarboxylation pathway. Preprint at 10.1101/2023.06.02.543384 (2023).

13. Folkes, L. K. & Wardman, P. Enhancing the efficacy of photodynamic cancer therapy by radicals from plant auxin (indole-3-acetic acid). Cancer Res 63, 776–779 (2003).

14. Fukuyama, T. T. & Moyed, H. S. Inhibition of Cell Growth by Photooxidation Products of Indole-3-acetic Acid. Journal of Biological Chemistry 239, 2392–2397 (1964).

15. Hayashi, K. et al. The main oxidative inactivation pathway of the plant hormone auxin. Nat Commun 12, 6752 (2021).

16. Hernandez-Garcia, J. & Weijers, D. The origin and evolution of auxin as a plant signaling molecule. Current Biology 36, R545–R552 (2026).

17. Jin, Q., Scherp, P., Heimann, K. & Hasenstein, K. H. Auxin and cytoskeletal organization in algae. Cell Biology International 32, 542–545 (2008).

18. Katz, E. et al. The glucosinolate breakdown product indole-3-carbinol acts as an auxin antagonist in roots of A rabidopsis thaliana. The Plant Journal 82, 547–555 (2015).

19. Klämbt, D., Knauth, B. & Dittmann, I. Auxin dependent growth of rhizoids of Chara globularis. Physiologia Plantarum 85, 537–540 (1992).

20. Kuhn, A. et al. RAF-like protein kinases mediate a deeply conserved, rapid auxin response. Cell 187, 130–148.e17 (2024).

21. Kurtović, K. et al. The role of indole-3-acetic acid and characterization of PIN transporters in complex streptophyte alga Chara braunii. New Phytologist 246, 1066–1083 (2025).

22. Kwiatkowska, M., Gosek, A. & Godlewski, M. Effect of GA3, IAA and their mixtures on the formation and development of cell systems in the vegetative and generative thallus of Chara vulgaris L. Acta Soc Bot Pol 60, 313–326 (2014).

23. Müller, K. et al. DIOXYGENASE FOR AUXIN OXIDATION 1 catalyzes the oxidation of IAA amino acid conjugates. Plant Physiology 187, 103–115 (2021).

24. Ohtaka, K., Hori, K., Kanno, Y., Seo, M. & Ohta, H. Primitive Auxin Response without TIR1 and Aux/IAA in the Charophyte Alga Klebsormidium nitens. Plant Physiol. 174, 1621–1632 (2017).

25. Papagiannakis, A., De Jonge, J. J., Zhang, Z. & Heinemann, M. Quantitative characterization of the auxin-inducible degron: a guide for dynamic protein depletion in single yeast cells. Sci Rep 7, 4704 (2017).

26. Paponov, I. A. et al. Auxin-Induced Plasma Membrane Depolarization Is Regulated by Auxin Transport and Not by AUXIN BINDING PROTEIN1. Front. Plant Sci. 9, 1953 (2019).

27. Prinsen, E. Auxin homeostasis: new roads in a tight network. Journal of Experimental Botany 76, 3260–3262 (2025).

28. Schmidt, V. et al. Phytohormone profiling in an evolutionary framework. Nat Commun 15, 3875 (2024).

29. Skokan, R. et al. PIN-driven auxin transport emerged early in streptophyte evolution. Nat. Plants 5, 1114–1119 (2019).

30. Stasinopoulos, T. C. & Hangarter, R. P. Preventing Photochemistry in Culture Media by Long-Pass Light Filters Alters Growth of Cultured Tissues. Plant Physiol. 93, 1365–1369 (1990).

31. Tuli, V. K. Reversal of cytokinin action by riboflavin during stage II micropropagation: The role of 3-methyleneoxindole. In Vitro Cell.Dev.Biol.-Plant 36, 532–536 (2000).

32. Vanneste, S., Pei, Y. & Friml, J. Mechanisms of auxin action in plant growth and development. Nat Rev Mol Cell Biol 26, 648–666 (2025).

33. Vosolsobě, S., Skokan, R. & Petrášek, J. The evolutionary origins of auxin transport: what we know and what we need to know. Journal of Experimental Botany 71, 3287–3295 (2020).

34. Wood, N. L. & Berliner, M. D. Effects of indoleacetic acid on the desmid Micrasterias thomasiana. Plant Science Letters 16, 285–289 (1979).

35. Zeng, H. Y. et al. Origin and evolution of auxin-mediated acid growth. Proc. Natl. Acad. Sci. U.S.A. 121, e2412493121 (2024).

